# Septins are involved at the early stages of macroautophagy in *S. cerevisiae*

**DOI:** 10.1101/043133

**Authors:** Gaurav Barve, Shreyas Sridhar, Amol Aher, Sunaina Singh, Lakshmeesha K.N., Ravi Manjithaya

## Abstract

Autophagy is a conserved cellular degradation pathway wherein a double membrane vesicle, called as an autophagosome captures longlived proteins, damaged or superfluous organelles and delivers to the lysosome for degradation^1^. We have identified a novel role for septins in autophagy. Septins are GTP-binding proteins that localize at the bud-neck and are involved in cytokinesis in budding yeast^2^. We show that septins under autophagy prevalent conditions are majorly localized to the cytoplasm in the form of punctate structures. Further, we report that septins not only localize to pre-autophagosomal structure (PAS) but also to autophagosomes in the form of punctate structures. Interestingly, septins also form small non-canonical rings around PAS during autophagy. Furthermore, we observed that in one of the septin Ts" mutant, *cdc10-5*, the anterograde trafficking of Atg9 was affected at the non-permissive temperature (NPT). All these results suggest a role of septins in early stages of autophagy during autophagosome formation.

---

In yeast, the nascent autophagosome (phagophore) formation site is known as the preautophagosomal structure or phagophore assembly site (PAS) and is perivacuolar located. Recent work has shown that the PAS is tethered to ER exit sites where multiple autophagy proteins localize and their hierarchical sequence of appearance has been determined^1,2^. The initiation of the autophagosome biogenesis involves the Atg1 complex (Atg1-Atg13-Atg17-Atg11-Atg29-Atg31) along with the Class III PI3 kinase complex (Vps34-Vps15-Vps30/Atg6-Atg14). The membrane source for the developing autophagosome is contributed by the trafficking of Atg9 along with its transport complex (Atg1-Atg11-Atg13-Atg23-Atg27-Atg2-Atg18-TRAPIII) to help build the initial cup-shaped structure, the phagophore^3–5^. Structural contribution for the formation of the autophagosome comes in the form of two scaffold-like proteins, Atg17 (required for general autophagy and is recruited at PAS in response to starvation conditions) and Atg11 (for selective autophagy and is recruited at PAS in nutrient-rich conditions). Structural and biochemical studies suggest that a dimer of Atg17-Atg31-Atg29 complex at one hand interacts with Atg1 through Atg13 and the other hand interacts with Atg9, forming a complex^6 7^ Atg1 with it’s EAT (Early Autophagy Targeting) domain interacts with peripheral membrane sources. Thus, the pentameric complex of Atg1-Atg13-Atg17-Atg31-Atg29 promotes initial clustering of Atg proteins followed by phagophore expansion^7^. Additional recruitment of the Atg5-Atg12-Atg16 complex as well as Atg8 allows the completion of the autophagosome^8^. Altering expression levels of Atg8 either by use of heterologous promoter or using a knockout of a negative regulator of Atg8, Ume6, have illustrated the role of Atg8 in the regulation of the size of the autophagosomes^9,10^. Apart from the above-mentioned Atg proteins that are involved in the early stages of autophagy, several other accessory proteins are reported. Arp2, for example, links actin and the autophagy machinery and is involved in the anterograde transport of Atg9 during autophagy in yeast^11^. Trs85, a component of TRAPIII complex directs guanine-nucleotide exchange factor (GEF) of Ypt1, a Rab GTPase, to the PAS and is involved in autophagosome formation^12^.

Septins are GTP-binding proteins and are mainly involved in the cytokinesis step of cell division in yeast^13^. There are five mitotic septins in yeast that consist of CDC10, CDC3, CDC12, CDC11 and SHS1 along with two meiotic septins SPR3 and SPR28. Where proteins such as Cdc10, Cdc3, Cdc12, and Cdc11 are crucial for the septin collar formation at the bud-neck junction, Shs1 has several auxiliary roles in its assembly and organization^14,15^. Septins have been implicated to function in the cell cycle, vesicular transport, membrane dynamics and other intracellular processes^16^. Septins are composed of several functional domains each of which has been shown to play distinct roles in various septin mediated processes. These domains also cooperate in septin oligomerization necessary for filament formation^17–19^. Septins have been implicated in autophagy in mammalian cells, where they form cage-like structures around the intracellular bacteria that co-localize with the autophagosome marker, LC3. It is believed that these structures entrap the bacteria, restrict their motility and target it for degradation by autophagy ^20,21^. Further insights regarding the association of septins with the autophagy machinery in capturing and degrading the bacteria remains to be studied. It is not known whether septins are involved in other autophagy-related pathways. Here we have identified septins (CDC10 and CDC11) in a screen for a subset of temperature sensitive (Ts^−^) mutants that are affected in selective autophagy of peroxisomes, pexophagy. We employed an immunoblotting based pexophagy assay that we successfully used for a previous screen^22^. The Pot1-GFP strains generated were subjected to the pexophagy assay at permissive (PT-22°C) and non-permissive temperature (NPT-37°C) in starvation conditions. During pexophagy, capture and degradation of Pot1-GFP containing peroxisomes result in accumulation of a free form of GFP in the vacuole with a concomitant decrease in Pot1-GFP levels. As expected the wild type cells showed an accumulation of free GFP over time while in cells lacking Atg1, a key kinase in the autophagy pathway, there was no degradation of Pot1-GFP at both the temperatures (Fig S1a). We found that the septin Ts^−^ mutants for CDC10 (*cdc10-1* and *cdc10-5*) and CDC11 (*cdc11-3*, *cdc11-4*, *cdc11-5*) showed considerable block in GFP release at NPT (Fig S1a). However, the other two septins, CDC3 (*cdc3-1* and *cdc3-3*) and cdc12 (*cdc12-1*) Ts^−^ mutants did not show such dramatic decrease (Data not shown). These results were also corroborated by a fluorescence microscopy based pexophagy assay (Fig S1b). Here delivery of GFP-labeled peroxisomes to the vacuole was identified as diffused GFP inside the vacuolar lumen. The number of cells showing free GFP inside the vacuole was significantly reduced when cells were incubated under the starvation condition at NPT than at PT in the above-mentioned Ts^−^ mutants (Fig S1b).

The core machinery for the formation and fusion of the autophagosomes is shared between the two pathways (pexophagy and autophagy) and, therefore, we tested septin Ts^−^ mutants for their involvement in general autophagy. Strains expressing GFP-Atg8, a marker for autophagosomes, were used in the GFP-Atg8 processing assay which is in principle similar to the pexophagy assay. Similar to the results obtained by pexophagy assays, we noticed considerable slowdown of autophagy flux in the cdc10 (*cdc10-1* and *cdc10-5*) and the cdc11 (*cdc11-3*, *cdc11-4*, *cdc11-* 5) Ts^−^ mutants (Fig S1c) but not in CDC3 (*cdc3-1* and *cdc3-3*) and CDC12 (*cdc12-1*) Ts^−^ mutants (Data not shown). We also observed that free GFP levels gradually increased over time in WT cells at NPT compared to PT. However in the case of Ts^−^ mutants the free GFP levels did not increase much over time. These results indicated that certain septin mutants affected autophagy and related pathways. To better understand and confirm the nature of mutations in the Ts^−^ mutants, we PCR amplified the mutant Ts^−^ genes and sequenced them (Figure S4 for the tabulation of the mutations). From the sequencing results, it was evident that all mutations were present in the GTP-binding domain and these mutations were previously shown to affect GTP binding and have a negative effect on oligomerization and filament formation^19,23^.

To follow up on these findings, we generated deletions in the 3 non-essential mitotic septins - CDC11, CDC10 and SHS1. The cell morphology, phenotype and growth conditions were consistent with previous reports^18,24^. The status of pexophagy and general autophagy was analyzed in these mutants and detected by both microscopy and western blot analysis. As the septin filaments are unstable at higher temperatures^18^, we carried out the autophagy assays at 22°C (PT) and 37°C (NPT). The CDC11 deletion mutant (*cdc11*Δ) displayed a complete block in pexophagy and general autophagy at NPT while in *cdc10*Δ and *shs1*Δ cells, pexophagy and general autophagy were affected considerably at NPT (Fig S2a and S2b). These results were also confirmed by microscopy where free GFP release inside the vacuole was observed only in a small population of cells which indicated pexophagy was affected (Fig S2c). Septins form hetero-octamers resulting from two tetramers interacting head-to-head, *i.e*., Cdc11-Cdc12-Cdc3-Cdc10-Cdc10-Cdc3-Cdc12-Cdc11^14,18^. Cdc3 and Cdc12 are part of the central core while Cdc11 and Cdc10 occupy the terminal ends. In the deletion strains, in spite of the absence of the terminal septin protein, the ability of septin assemblies to interact is not wholly compromised at PT. Cdc3 and Cdc12 have the intrinsic ability as all septins to self-dimerise, although with reduced capability in the absence of Cdc10 and Cdc11 respectively. In the absence of Cdc10 or Cdc11, hetero-hexamers are formed which subsequently form partial higher-oligomers that are destabilized at higher temperatures^18^. This explains why the deletion strains at a lower temperature (PT) show autophagic activity but at elevated temperatures (NPT) are unable to do so.

Shs1 has the ability to occupy the position of Cdc11. In the wild-type cells, it functions to cap the ends and regulate the septin ring assembly^18,25^. The synergistic effect of the weak interaction of Cdc12-Cdc12 in the absence of Cdc11 and the increased functionality of Shs1 therein at higher temperatures might be resulting in premature termination of growth towards filament formation leading to a complete block in autophagy in *cdc11*Δ strain^24^. A similar scenario might be unfolding with Cdc10 but like Cdc11, it does not have an antagonistic analogue thereby allowing Cdc3 to retain dimerization and subsequently participate in autophagic flux although at reduced rates. To test if oligomerization property of septin assemblies is the key for its role in autophagy, we deleted SHS1 in a CDC11 null background. Interestingly, in the absence of the negative antagonist of Cdc11, autophagy was partially recovered in these double deletion strains (Fig S2a and S2b panel 6 from top). These results point at a cellular scenario where the higherorder oligomeric structures of the septin proteins are critical for the autophagic process.

Next, we wanted to study the effect of various domain deletion mutants of septins on pexophagy. Binding of GTP to the GTP-binding (G) domain plays a crucial role as it induces a conformational change in the septin structure that influences both the GTP-binding (G) interface as well as amino and carboxy (NC) terminal structural interfaces. Septin mutants having mutations in the G domain failed to form the cortical ring at the bud-neck^26,27^. The two interfaces are essential for septin-septin interaction, subsequent oligomerization and filament formation. The ability of septins to hydrolyze GTP is still highly debatable ^23,28–30^. To strengthen our claim that structural conformations of septins are required for their role in autophagy, we analyzed point mutations in the GTP-binding domain and other domain deletions mutations involved in septin-septin interactions. Here we focused on Cdc11p as the block in autophagy observed in its absence was nearly absolute. GTP binding mutants having mutations in the P-loop (*cdc11-Δ18-20*) and the GTP binding domains (*cdc11-G29A*, *G32A*, *G34A*, *cdc11-R35A* and *cdc11-G230E*) exhibited partial to severe effect on selective and general autophagy (Fig S3a and S3b). These mutations have been shown to be defective in GTP binding resulting in reduced septin-septin interaction^17^. Among the domain deletion mutants, autophagy was also defective at both PT and NPT in the coiled-coil (CC) domain deletion mutant, *cdc11*-Δ347-415 (Fig S3b). This domain is necessary for the formation of the NC interface and in turn critical for septin-septin interaction. The coiled-coil domain mutant lacks in its ability to form short oligomers of the septin tetramer *via* Cdc11 dimerization. Another domain mutant that was blocked in autophagy at NPT had a deleted PI (phosphatidylinositol)-binding domain (residues 18-20 amino acids). Yet mutants that had the basic residues mutated (R12Q, K13Q, R14Q, K15Q, H16Q) in the PI binding region, did not show any significant block in autophagy (Data not shown). In these mutants, the possibility that the domain deletions are affecting the structural conformation of septins resulting in the observed phenotype, cannot be ruled out (Fig S3b). These mutants further strengthen the possibility of a structural role for septins in autophagy. Although the ring forming capability at the bud-neck region is not critical, the ability of septin octamer or tetramer to oligomerize into high-order elements and possibly filaments is essential.

To understand the context of septin protein localization during starvation conditions, we examined cells containing GFP-tagged Cdc10p, Cdc11p and Shs1p individually. As expected, in growth conditions all three tagged mitotic septins localized to the bud-neck region forming septin collar in dividing cells (Fig 1a, starvation 0h). On subsequent transfer of cells into starvation media, the septin proteins were seen at the cell periphery forming distinct puncta (Fig 1a, starvation 2h, 4h and 24h). To confirm our observation, we treated the cells with rapamycin, an inducer of autophagy, whilst growing them in rich media. Like the starvation media, punctate structures were observed on treatment with rapamycin as compared to rich media. Upon quantitation, it was evident that under starvation and rapamycin treated conditions, the number of cells showing punctate structures was greater than the number of cells showing ring at the budneck (Fig 1b). These results with tagged septins reveal a pattern of re-localization from the budneck to the cell periphery when faced with starvation stress. Because during starvation conditions, wild-type septins showed punctate structures and septin Ts^−^ mutants also appeared as single punctum (data not shown), it was tempting to speculate that the perivacuolar localized dotlike structure could be an autophagosome organizing center, the pre-autophagosomal structure (PAS). We subsequently asked if these punctate structures in the wild-type cells were localized to any known autophagic structures. To substantiate their role in autophagy, we created strains with GFP-labeled septin and mCherry labeled Atg8 (a PAS and an autophagosomal marker). In the wild-type background, these cells were microscopically examined under nutrient rich and starvation conditions. We observed one-third of the cells showing co-localization between septins (Cdc10, Cdc11 and Shs1) and Atg8 at the PAS near vacuole (Fig 1c and 1d). When the images were meticulously observed we found that in starvation conditions, apart from the punctate structures, septins formed ring-like structures which were different from the canonical rings in localization and in size (Fig 2a, 2b and 2c). Unlike the canonical rings that are seen at the bud-neck region, these non-canonical rings were rarely observed and were localized to regions other than the bud-neck region and were comparatively smaller in size (400-600nm). Surprisingly, these non-canonical rings also co-localized at the PAS (Fig 2a, 2b, 2c and Supplementary video 1). From the analysis of the 3D images, Shs1-GFP cells formed a non-canonical ring around the PAS at initial Z planes and it co-localized with PAS at later Z planes suggesting completion of ring formation (Fig 2b and Supplementary video 2). Furthermore, as shown in figure 2b, in later Z sections we also observed a vesicle-like structure of ~600nm diameter which has a similar diameter to that of the reported diameter of an autophagosome^31^. Similar to Shs1-GFP, Cdc10-GFP cells also showed the formation of the non-canonical ring which co-localized with mcherry-Atg8 (mch-Atg8) and subsequently formed a vesicle-like structure of ~600nm size (Fig 2c and Supplementary video 3). Thus, the results so far suggest that septins might be playing an early role in autophagosome formation most likely by providing a scaffolding platform for the developing autophagosome. In order to know whether septins are associated with only PAS or they are associated with autophagosomes also, we checked the colocalization of Cdc10-GFP and Shs1-GFP with mch-Atg8 in the *ypt7*Δ background. Ypt7 is a Rab GTPase which plays a major role in autophagosome-vacuole fusion^32,33^ and hence in the absence of Ypt7, autophagosomes are accumulated inside the cytoplasm. Surprisingly, we found that Cdc10-GFP, Cdc11-GFP and Shs1-GFP puncta co-localized with mch-Atg8. We observed that more than one mch-Atg8 punctum co-localized with Cdc10-GFP, Cdc11-GFP and Shs1-GFP puncta (Fig 3a and 3b). Similar to wild-type cells (Fig 2), we also observed non-canonical ring formation by Cdc10-GFP and Shs1-GFP around mch-Atg8 in *ypt7*Δ cells (Fig 3c and 3d). Thus, the single co-localization event at PAS in the case of wild-type strains and multiple colocalization events in *ypt7*Δ strains suggested that septins are associated not only with PAS but also with the autophagosomes.

**Fig. 1.**
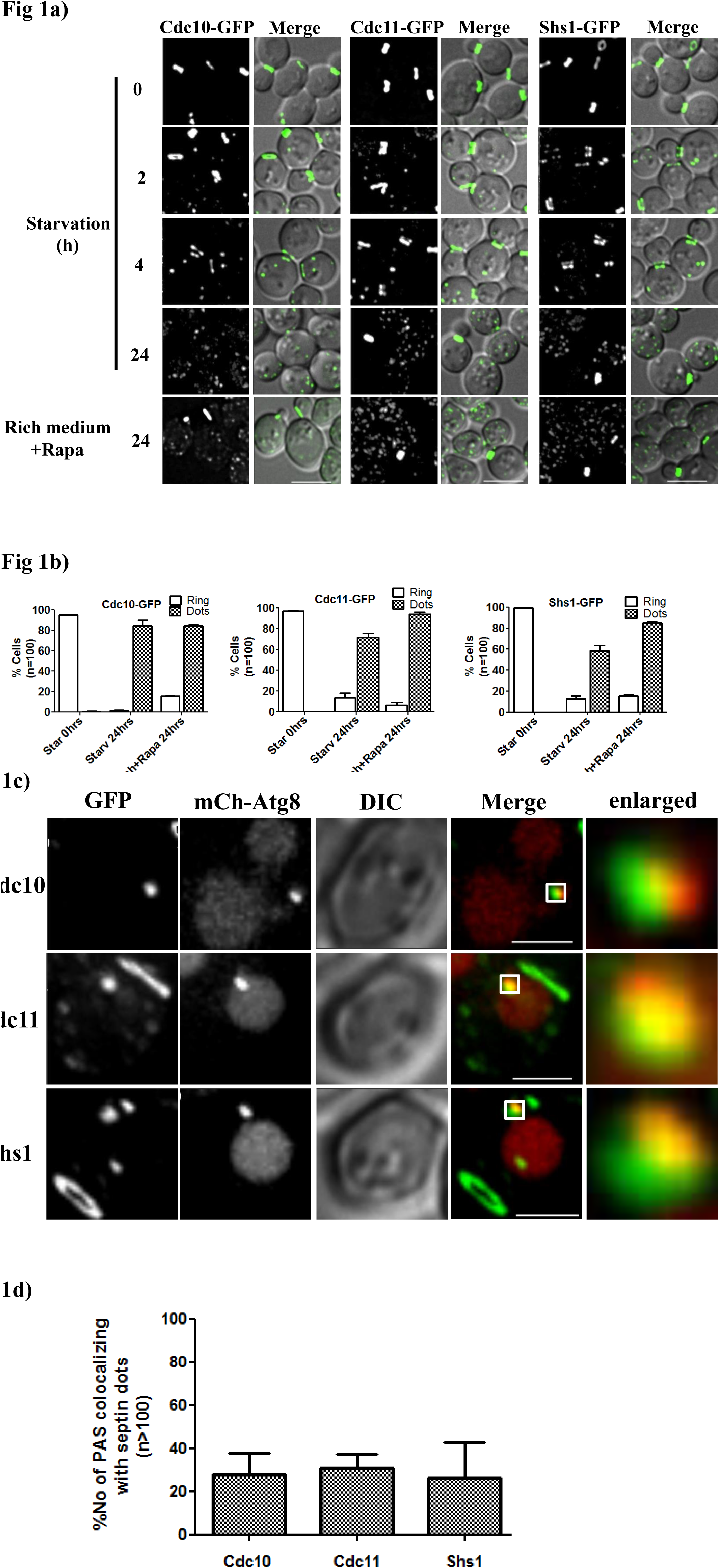
Septins formed punctate structure during starvation and were localized at PAS. **a)** Microscopy images of Cdc10-GFP, Cdc11-GFP and Shs1-GFP under nutrient rich, nutrient deficient and rapamycin treated conditions. Cells from log phase (0.6 to 0.8 A600) were transferred to starvation medium and imaged at different time points. **b)** Quantitation of the number of cells showing rings and puncta grown in rich and starvation media for 24hr. For quantitation, cells showing only ring or only dots were considered. Images acquired were converted to maximum intensity projections, de-convoluted and a total of 100 cells were quantitated. Scale bar 5μm. **c)** Septins co-localize with mcherry-Atg8, a PAS and an autophagosome marker. Cells growing in log phase were incubated in starvation medium for 4hr in nitrogen deficient media and then imaged. All images are single Z-stack and are deconvoluted using SoftWorx software (Delta vision). **d)** Quantitation of the number of PAS that co-localize with septins. Quantitation was done manually using cell counter plugin of Fiji at every z section.

**Fig. 2.**
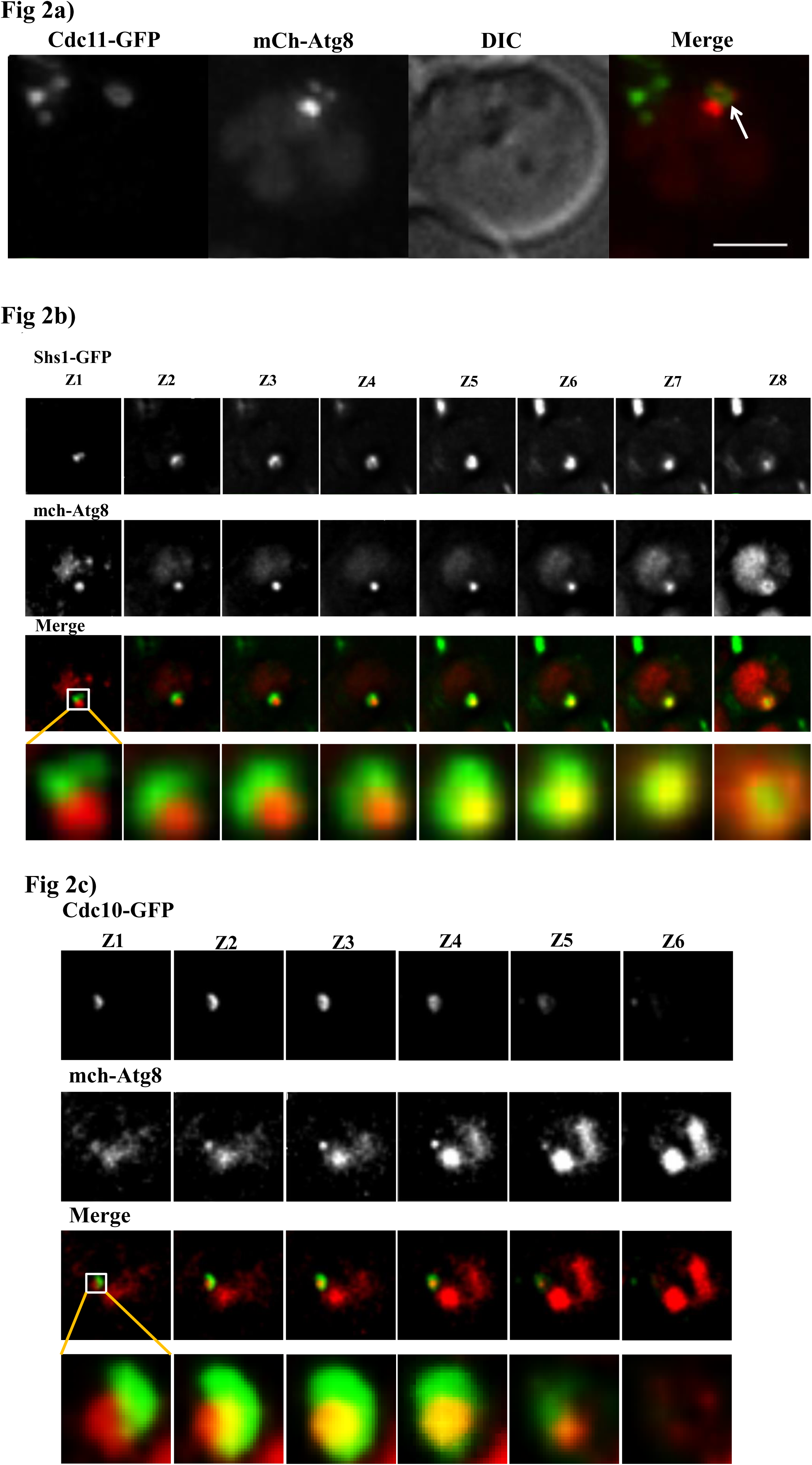
Septins formed a non-canonical ring-like structures around PAS during starvation. **a)** Formation of the non-canonical ring (white arrow) by Cdc11 that co-localize with PAS. **b)** and **c)** Formation of a non-canonical ring around mcherry-Atg8 by Shs1 and Cdc10 respectively. Shs1-GFP and Cdc10-GFP Cells were incubated in starvation medium for 4hr and were imaged. Z-stacks of 200nm each is shown. Scale bar 1μm.

**Fig. 3.**
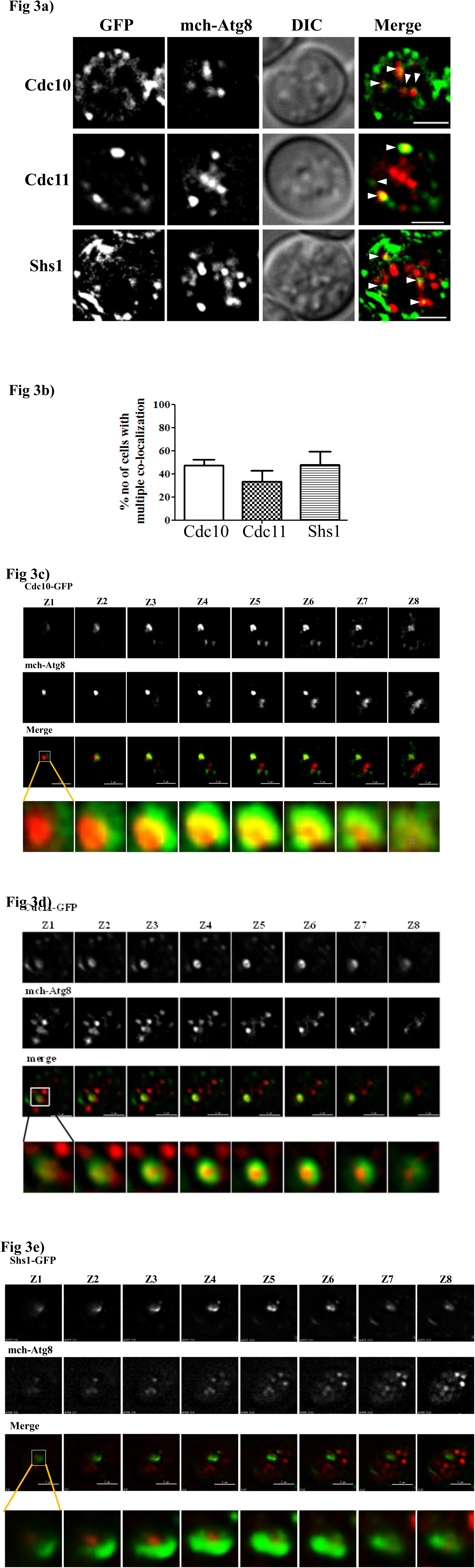
Septins localized at autophagosomes during starvation. **a)** Septins co-localize not only at PAS but also at autophagosomes which are shown by white arrow heads. Cdc10-GFP *ypt7*Δ, Cdc11-GFP *ypt7*Δ and Shs1-GFP *ypt7*Δ cells expressing mcherryAtg8 were incubated in starvation medium for 4hr and were imaged. **b)** Quantification of the number of cells showing multiple co-localizations with mCherry-Atg8 puncta. For quantitation images were deconvolved, background subtracted and then co-localization was checked using co-localization highlighter plugin and co-localized points were then quantitated using cell counter plugin of Fiji. More than 150 cells were quantitated and the graph shows the mean of three independent experiments with standard error. **c)** Non-canonical ring formation around mcherry-Atg8 by Cdc10GFP d) Non-canonical ring formation around mcherry-Atg8 by Cdc11-GFP cells. **e)** Non-canonical ring formation around mcherry-Atg8 by Shs1-GFP cells.

Similar to septins, Atg9, an important autophagy protein also shows multiple puncta during starvation^34^. This observation tempted us to check whether septins co-localize at Atg9 vesicles as well. Thus, we created Septin-GFP and mcherry-Atg9 strains and imaged them. Interestingly, one of the puncta of Cdc10-GFP, Cdc11-GFP and Shs1-GFP co-localized with one of the mchAtg9 puncta after 4h of starvation (Fig4a and 4b). During early stages of autophagosome biogenesis, Atg9 trafficking from peripheral membranes reservoirs such as the ER and mitochondria brings membrane to the developing phagophore^35–37^. From peripheral locations, Atg9 mobilizes to the PAS during starvation bringing membrane required for an autophagosome formation (anterograde trafficking) and is then trafficked away from PAS with the help of Atg1 complex (retrograde trafficking)^5,38,39^. We checked Atg9 trafficking during starvation in the septin Ts^−^ mutants and in the septin deletion strains. Unlike all other septin mutants (data not shown), we observed that *cdc10-5* strain showed single bright Atg9 punctum with few less bright

Atg9 puncta present at the peripheral sources at NPT which was similar to *atg1*Δ strain^5^ (Fig 5a and 5b). Further, we checked co-localization of the bright Atg9 punctum with the PAS marker ApeI-RFP. Surprisingly, in *cdc10-5* strain, the co-localization was decreased almost by 50% as compared to *atg1Δ* strain, implying accumulation of Atg9 vesicles at a place other than PAS and hence suggesting that an anterograde Atg9 transport is affected in *cdc10-5* strain (Fig 5c and 5d). The *cdc10-5* strain has a point mutation (G44D) in GTP binding domain and GTP binding has been shown to be important for septin filament and hence higher order structure formation^19,29^. Thus, all these observations highlight the role of GTP binding and septin filament formation in the anterograde transport of Atg9.

**Fig. 4.**
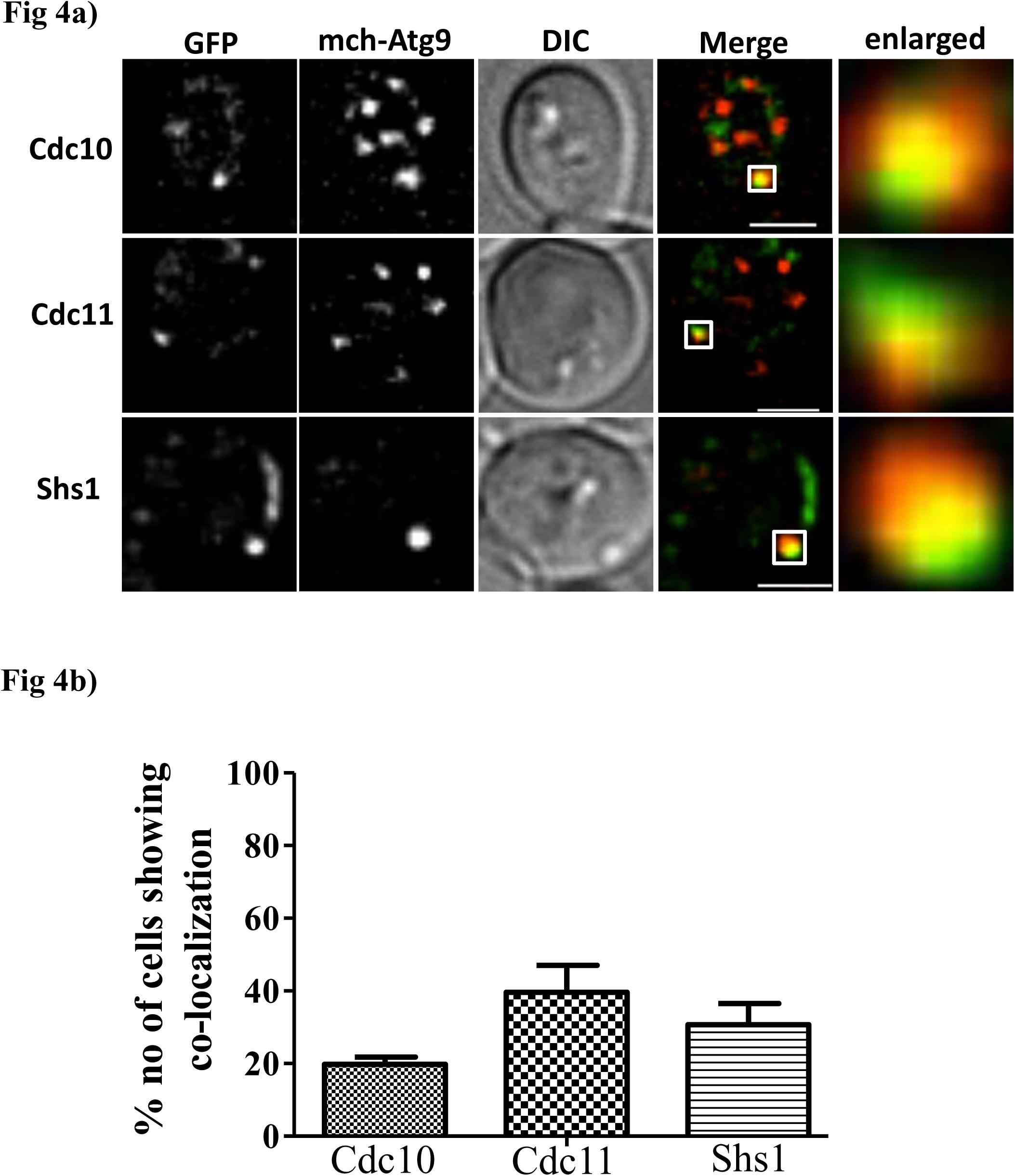
Septins partially co-localized with Atg9 during starvation. **a)** Septins co-localize with mcherry-Atg9. Cells were grown as mentioned in Fig 2c) and were imaged. **b)** Quantification of the number of cells showing co-localization with mcherry-Atg9 puncta. Quantitation was performed as mentioned in 3 b). More than l50 cells were quantitated and the graph shows the mean of three independent experiments with standard error.

**Fig. 5.**
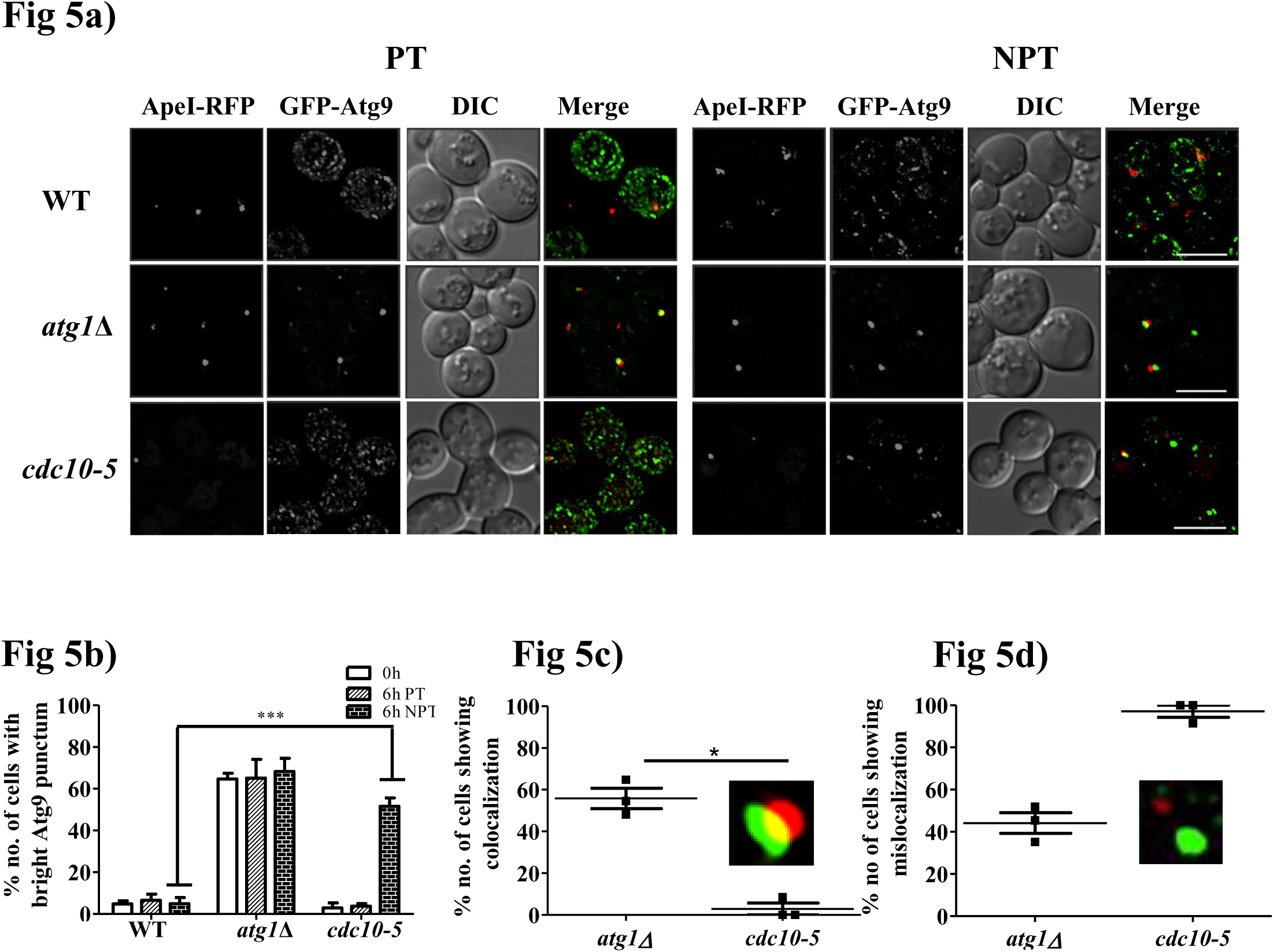
Cdc10 is involved in the anterograde transport of Atg9. a) Single bright Atg9 punctum formation in *cdc10-5* at NPT. Cells were grown in starvation medium for 6hr and were imaged. All images are maximum intensity projections (MIP) and are deconvoluted using SoftWorx software (Delta vision). Scale bar 5 μm. b) Quantitation of the number of cells showing bright Atg9 punctum at PT and NPT. Quantitation was done manually using Fiji and total of 50 cells were quantitated in each of the three experiments and mean of the three experiments with the standard error was plotted, p<0.00l, PT versus NPT in Wild-type and in *cdc10-5* cells. c) Quantitation of the number of cells showing co-localization between bright Atg9-GFP punctum and PAS marker ApeI-RFP. d) Quantitation of the number of cells showing mislocalization between bright Atg9-GFP punctum and ApeI-RFP. Quantitation was done manually using Fiji at each z section and a total of 30 cells was quantitated in each of the three experiments. The mean of the three experiments with the standard error was plotted, p<0.05, *atg1*Δ versus *cdc10-5* cells.

In summary, we have found a novel role of septins in autophagy. Compared to wild-type cells, septin mutants showed significantly affected phenotype in general autophagy as well as pexophagy at NPT. Proper septin rod and subsequent filament formation might be important for autophagy function as at NPT both filament assembly^18^ and autophagy are blocked in septin mutants. All the three wild-type septins *i.e*. Cdc10, Cdc11 and Shs1 localized to the cytoplasm as puncta upon nitrogen starvation. Septins also localized at PAS and showed non-canonical ring formation around PAS. Further septins were also found to associate with autophagosomes but did not go along with autophagosomes to the vacuole. This implies a role of septins at an early stage of either PAS organization or autophagosome formation. Since in Cdc10-GFP, Cdc11-GFP and Shs1-GFP strains, free GFP was not observed in the vacuole under starvation, it is highly unlikely that septins are degraded by autophagy. Furthermore, our data suggests an additional role of septins in helping Atg9 molecules to deliver membrane source for developing autophagosomes. In conclusion, septins associate with various steps of autophagosome biogenesis.

## Materials and methods

### Yeast Strains and media

Yeast wild type and autophagy knockout mutant strains used in this study are BY4741, BY4742, and S288C and were obtained from EUROSCARF. Strain, plasmid and primers details are given in the supplementary table T1. Wild-type Pot1-GFP and septin mutant strains are laboratory strains with genomically tagged GFP to the C-terminus of Potl (HIS selection marker), obtained from Prof. Rachubinski, University of Alberta, Canada. Septin Ts^−^ mutants were kindly provided by Prof. Charlie Boone, Toronto and septin knockout mutants were prepared using the standard transformation protocols^40^. GFP-ATG8 pRS316 and mcherry-ATG8 pRS316 plasmids were a kind gift from Prof. Ohsumi, Tokyo. ATG9-GFP pRN295 was a gift from Prof. Michael Thumm, Stuttgart, Germany.

Wild-type cells and mutants were grown in YPD medium (1% yeast extract + 2% peptone + 2% dextrose) at 30°C and 22°C respectively. For pexophagy assays oleate medium (0.25% yeast extract, 0.5% peptone, l% oleate + 5% Tween-40 and 5 mM phosphate buffer) was used to induce peroxisome formation and synthetic defined medium (0.17% yeast nitrogen base lacking amino acids and ammonium sulphate + 2% dextrose) was used to induce autophagy. For Ts^−^ mutants and knockout mutants 22°C and 37°C were used as permissive and non-permissive temperatures respectively.

### Microscopy and cytology

Cells were grown in respective media, centrifuged and were mounted on agarose (2% w/v) pad for microscopy. Images were taken in Z sections of 0.2μm step size using Delta Vision microscope (Applied Precision) fitted with 100x l.4NA objective and Cool-SNAP HQ2 camera. Image processing and quantitation were performed using SoftWorx (Applied Precision) and Fiji (NIH). For co-localization analysis, images were de-convolved; background subtracted and colocalized entities were either quantitated manually using cell counter plugin or automatically using co-localization highlighter plugin in Fiji at all Z sections.

### Western blot analysis

Whole cell extracts were prepared by TCA (12.5% TCA w/v) precipitation method followed by ice-cold acetone washes (twice) and lysates were stored at −80°C before the actual analysis. Protein extracts were then analyzed using SDS-PAGE and Western blot (mouse anti-GFP monoclonal 1:3000, Roche applied Science). Blots were visualized using anti-mouse secondary antibody conjugated to HRP (BIORAD) (1:10000) on a gel documentation system (G: Box chemi XT4, Syngene).

### Statistics

All statistical analysis was performed using Prism and to calculate significance levels two-way ANOVA and Student’s t-test were used.

## Acknowledgements

We would like to thank Prof. Charles Boone, Prof. Yoshinori Ohsumi, Prof. Kausik Chakraborthy, Prof. Michael Snyder, Prof. Michael Thumm, and Prof. Richard Rachubinski for generously sharing strains, plasmids and reagents. Critical reading of the manuscript and inputs from Prof. MRS Rao, Prof. Subba Rao, Aparna Hebbar and members of the autophagy lab is appreciated. This work was supported by Wellcome Trust/DBT India Alliance Intermediate Fellowship (509159/Z/09/Z) and JNCASR intramural funds to RM.

